# Assessing Immune Microenvironment in TCGA-LUAD via CIBERSORTx Using Single-Cell Derived Signature Matrix and ESTIMATE Algorithm

**DOI:** 10.1101/2024.05.08.592760

**Authors:** Madhulika Verma

## Abstract

Lung cancer (LC) remains a significant global health concern, affecting millions worldwide each year. Tumor-infiltrating immune cells (TIICs) play a crucial role in Lung Cancer progression and prognosis, with various immune cell types infiltrating the tumor microenvironment. Traditional methods like immunohistochemistry and flow cytometry have limitations in accurately profiling TIIC subtypes. However, recent advancements in single-cell RNA sequencing and computational algorithms like CIBERSORTx offer a promising approach for characterizing TIICs in bulk tumor samples.

In this study, we undertook the validation of the signature matrix comprising 14 distinct immune cell types and subtypes, which was originally derived from PBMC single-cell RNA-seq data, in our previous work (Verma, 2024). The positive controls included 8 bulk RNA-seq samples of whole blood and specific immune cell bulk RNA-seq samples, while the negative control comprised neuroblastoma cell lines lacking immune content. Subsequently, we applied this signature matrix to deconvolute TCGA-LUAD data (n = 598), and assessed tumor purity and immune-stromal content using the ESTIMATE algorithm.

Our findings indicate that the signature matrix accurately reflected flow cytometry-derived fractions, supported by correlation analysis. Specifically, the second positive control and negative control accurately reflected immune and non-immune sample fractions, respectively, further validating the efficacy of our approach. This study also provide insights into the invasion of immunocytes in lung adenocarcinoma and highlight the potential of computational tools like CIBERSORTx and ESTIMATE in characterizing the immune microenvironment of LC.

## 1. Introduction

Each year, an alarming 2.3 million individuals worldwide face the daunting diagnosis of lung cancer, with over 200,000 cases reported in the USA alone (Lung Cancer - Non-Small Cell - Introduction, 2012). While cigarette smoking stands as the primary culprit, it’s essential to acknowledge that lung cancer can afflict anyone, irrespective of smoking history. Moreover, lung cancer, regardless of its extent or spread, offers reasonable prospects for treatment.

The stigma associated with smoking often clouds perceptions of lung cancer patients, potentially hindering access to support and care (Lung Cancer - Non-Small Cell - Introduction, 2012). However, it’s critical to dispel the misconception that only smokers are at risk. In reality, a significant proportion of lung cancer cases occur in individuals who have never smoked or ceased smoking years prior (Lung Cancer - Non-Small Cell - Introduction, 2012).

The lungs boast a rich tapestry of cellular diversity, primarily comprising epithelial cells responsible for mucus secretion, vital for lung hydration and defense. Complementing these are hormone-producing cells, structural support elements, nerve cells, and blood cells, collectively ensuring optimal lung function (Lung Cancer - Non-Small Cell - Introduction, 2012).

Lung cancer, comprising small-cell lung cancer and non-small cell lung cancer (NSCLC), presents a diverse spectrum of malignancies warranting distinct treatment modalities. NSCLC, originating from normal lung cells undergoing metastatic transformation, manifests as neoplasms within the lung tissue, ranging from benign to malignant tumors. Notably, these malignant tumors can disseminate cancerous cells via blood or lymphatic circulation, often targeting the chest’s central region as a primary site of spread (Lung Cancer - Non-Small Cell - Introduction, 2012).

NSCLC encompasses several subtypes, each with distinct histological features and clinical implications. Adenocarcinoma, the most prevalent NSCLC subtype, arises from lung periphery epithelial cells and accounts for nearly 40% of NSCLC cases. Conversely, squamous cell carcinoma originates from specialized planar cells lining the lung’s inner surface, representing approximately 30% of NSCLC instances. Large cell carcinoma, although less common, exhibits striking morphological differences and is often categorized under adenocarcinoma or squamous cell carcinoma due to diagnostic advancements (Lung Cancer - Non-Small Cell - Introduction, 2012).

In the context of lung cancer (LC), the infiltration of tumor-infiltrating immune cells (TIICs) plays a pivotal role in prognosis and treatment outcomes (Meza et al., 2015). LC, being immunologically sensitive, attracts a heterogeneous array of TIICs, including T lymphocytes, dendritic cells (DCs), macrophages, mast cells, and neutrophils. The abundance, composition, and localization of tumor-infiltrating lymphocytes (TILs) serve as crucial prognostic indicators (Gentles et al., 2015). Previous methodologies such as immunohistochemistry and flow cytometry, while informative, presented limitations in capturing the full spectrum of immune populations. Leveraging gene expression data and the CIBERSORTx (Steen et al., 2020) algorithm, researchers can extrapolate estimates of cellular subtypes within tumor samples. Utilizing a “signature matrix,” a bespoke expression profile of cell type-specific marker genes, CIBERSORTx facilitates precise deconvolution, offering insights into the immune landscape of bulk tumor samples (Steen et al., 2020).

In this work, we first used the positive and negative control bulk RNA-seq samples to validate the custom signature matrix of 14 different immune cell types and subtypes made from PBMC single-cell RNA-seq data, constructed in the previous paper (Verma, 2024). We then proceeded with this custom signature matrix to perform the deconvolution of TCGA-LUAD (*The Cancer Genome Atlas Program*, n.d., *Home* | *NCI Genomic Data Commons*, n.d.) data (lung adenocarcinoma, n = 598) to assess the invasion of immunocytes in the tumor. We also used ESTIMATE (Yoshihara et al., 2013)(Estimation of Stromal and Immune Cells in Malignant Tumor Tissues Using Expression Data), which uses inbuilt gene expression signatures to evaluate the purity of the tumor and assess the overall immune and stromal content of the LUAD. ESTIMATE works on the principle of ssGSEA (single sample gene set enrichment analysis) (Barbie et al., 2009).

## 2. Materials and Methods

### 2.1. Data procurement

#### Validation data

For validation of the custom signature matrix, we acquired three types of publicly available data: two positive controls and one negative control. The “Gene Expression Omnibus” (Edgar, 2002, Barrett et al., 2012) was used to get all of the data in the form of FPKM and TMM values.

The first positive control was **GSE127813** (Newman et al., 2019), which had whole blood samples collected from 12 healthy adults. The leucocyte composition was immediately enumerated after extraction via FACS. We had the FACS-derived composition of T cells, Neutrophils, Monocytes, CD4 T lymphocytes, CD8 T lymphocytes, B cells, total Lymphocytes, and NK cells. The Lymphocytes + Monocytes + Neutrophils = 1. This means that the total myeloid content was monocyte, and the total lymphoid content (CD4 T lymphocytes, CD8 T lymphocytes, B lymphocytes, and NK cells) was lymphocyte. Similarly, T lymphocytes = CD4 T lymphocytes + CD8 T lymphocytes. This relative configuration was necessary while comparing the CIBERSORTx predicted fractions.

The second positive control was **GSE60424** (Linsley et al., 2014), which had whole transcriptome profiling of whole blood and the immune cell subsets. These samples were taken from healthy and diseased patients (multiple sclerosis, amyotrophic lateral sclerosis, sepsis, and type 1 diabetes) before and after 24 hours of the first IFN-beta treatment. The immune cell subsets extracted were precisely isolated groups of neutrophils, monocytes, B lymphocytes, CD4 T lymphocytes, CD8 T lymphocytes, and natural killer cells. There were 134 samples-20 whole blood, 20 monocytes, 14 NK cells, 20 B cells, 20 CD4 T cells, 20 CD8 T cells, and 20 neutrophils.

The negative control was **GSE89413** (Harenza et al., 2017), which had whole transcriptome profiling of 39 commonly used neuroblastoma cell lines. These cell lines had varying genomic characteristics, disease stages, and phases of therapy. It also had hTERT-immortalized retinal pigmented epithelial cell line (n = 1) and neural cell RNA from the fetal brain (healthy, n = 1).

#### TCGA-LUAD data

The “TCGAbiolinks” R package was used to get lung adenocarcinoma datasets from the TCGA (Colaprico et al., 2015). The category of the data was “Transcriptome Profiling”, the type of the data was “Gene Expression Quantification”, and the analysis type was “STAR - Counts”. These were the technical terms that were used to extract the desired data. The data consisted of 60,660 genes from 598 patients (samples). The other specifications for this data were as follows:

- Out of 598 samples:
  - Primary Tumor - 537
  - Recurrent Tumor - 2
  - Solid Tissue Normal - 59

- Out of 598, the vital status:
  - Alive - 380
  - Dead - 218

- Out of 598, pathologic_stage
  - Stage I - 5
  - Stage IA - 152
  - Stage IB - 169
  - Stage II - 1
  - Stage IIA - 55
  - Stage IIB - 82
  - Stage IIIA - 85
  - Stage IIIB - 12
  - Stage IV - 28

- Primary Diagnosis (Histological subtypes)
  - Acinar cell carcinoma - 22
  - Adenocarcinoma with mixed subtypes - 114
  - Adenocarcinoma, NOS - 374
  - Bronchio-alveolar carcinoma, mucinous - 5
  - Bronchiolo-alveolar adenocarcinoma, NOS – 3
  - Bronchiolo-alveolar carcinoma, non-mucinous - 22
  - Clear cell adenocarcinoma, NOS - 3
  - Micropapillary carcinoma, NOS - 3
  - Mucinous adenocarcinoma - 20
  - Papillary adenocarcinoma, NOS - 25
  - Signet ring cell carcinoma - 1
  - Solid carcinoma, NOS - 6

In addition to these details, the data had full clinical details, such as whether or not the person smoked, the molecular subtype, the type of treatment, the person’s age, gender, race, and many more.

### 2.2. Merging the custom signature matrix

Since our validation data only had six major immune subsets, we had to turn our custom signature matrix of 14 immune subsets into six major ones. By taking the average of each gene’s value in the custom signature matrix, the following is how the signature matrix was merged:

- All B cell subsets (Activated/matured B cells and memory B cells) were merged together to form B cells,
- All effector, memory, cytotoxic and naive CD8 T cells were merged together to form the CD8 group.
- Naive CD4 were made CD4 T cells
- All the myeloids (plasmacytoid DC, Monocyte DC, CD14 Monocytes, CD16 monocytes, and macrophages) were merged together resulting in the Monocyte group.
- Both NK cell groups (NK cells and Activated NK cells) were merged into one, forming the NK cell group
- Neutrophils were kept as they were.

### 2.3. Deconvolution of Validation Data and TCGA-LUAD Data

#### Validation data

The “impute cell fractions” utility of CIBERSORTx was used to “deconvolve” both the positive and negative controls. Because there are technical differences between various platforms and tissue-preservation methods (such as “FFPE vs. fresh-frozen tissues”), CIBERSORTx offers a correction method for dealing with the platform-induced errors, that makes it feasible for the implementation of a signature matrix to bulk data of GEPs in spite both being obtained from different protocols. There were two modes for batch correction in CIBERSORTx: bulk (B-mode) and single-cell (S-mode). The S-mode batch correction was turned on since our signature matrix was from the single-cell platform, and the quantile normalization was turned off, as suggested by the CIBERSORTx assistant team for RNA-seq data being the mixture. All three validation datasets had relative mode turned on, and the number of permutations was kept at 100. In addition to this run, the positive control 2 and the negative control were also run in “absolute” mode with the rest of the parameters kept the same to compare the total immune content present in the positive and negative samples. In order to ensure that the total fractions for a particular mixing sample equal 1, by convention, CIBERSORTx calculates the relative proportion of each cell type within the signature matrix. The overall abundance of each cell type gets quantitatively measured by the absolute score. In a nutshell, the absolute score is “the median expression level of every gene in the signature matrix divided by the median expression level of every gene in the mixture.” (Chen et al., 2018)

#### TCGA-LUAD data

After the deconvolution of the validation datasets, the TCGA-LUAD data was processed to convert the Ensembl gene IDs to the gene symbols, and the redundant genes were removed. After the pre-processing, the data consisted of 59,427 genes from 598 samples. Using the “impute cell fractions” tool on the CIBERSORTx portal, the “deconvolution” was then performed on this data. Here, the original custom signature matrix of 14 immune cell types was used. S-mode batch correction was enabled, and quantile normalization was disabled. The permutations were kept at 100. The run was made with the “absolute” mode turned on.

### 2.4. Statistical analysis

R was used to look at the relationship between the flow cytometry and the CIBERSORTx predicted fractions of immune cells in the 12 healthy whole blood samples from “positive control 1.” The plots were generated in R using the package “ggplot2” (Wickham, 2016).

### 2.5. ESTIMATE analysis

The ESTIMATE analysis was performed in R on the TCGA-LUAD data to check the degree of immune and stromal infiltration in the cancer data and also to calculate the purity of the tumor.

## 3. Results and Discussion

### 3.1. Validation data

#### Positive control 1

Deconvolution was done on the 12 whole blood samples using the merged Custom signature matrix of the six major cell types. As a result, the relative fractions for each immune subset were supported by metrics like the p-value, Pearson correlation coefficient, and RMSE. The figure 1. shows the relative fractions of the cell types per sample. It should be noted that the immune cell fractions derived from flow cytometry were also reported in relative fractions per sample, so the lymphoid + myeloid + neutrophil content summed up to one.

**Figure 1.**
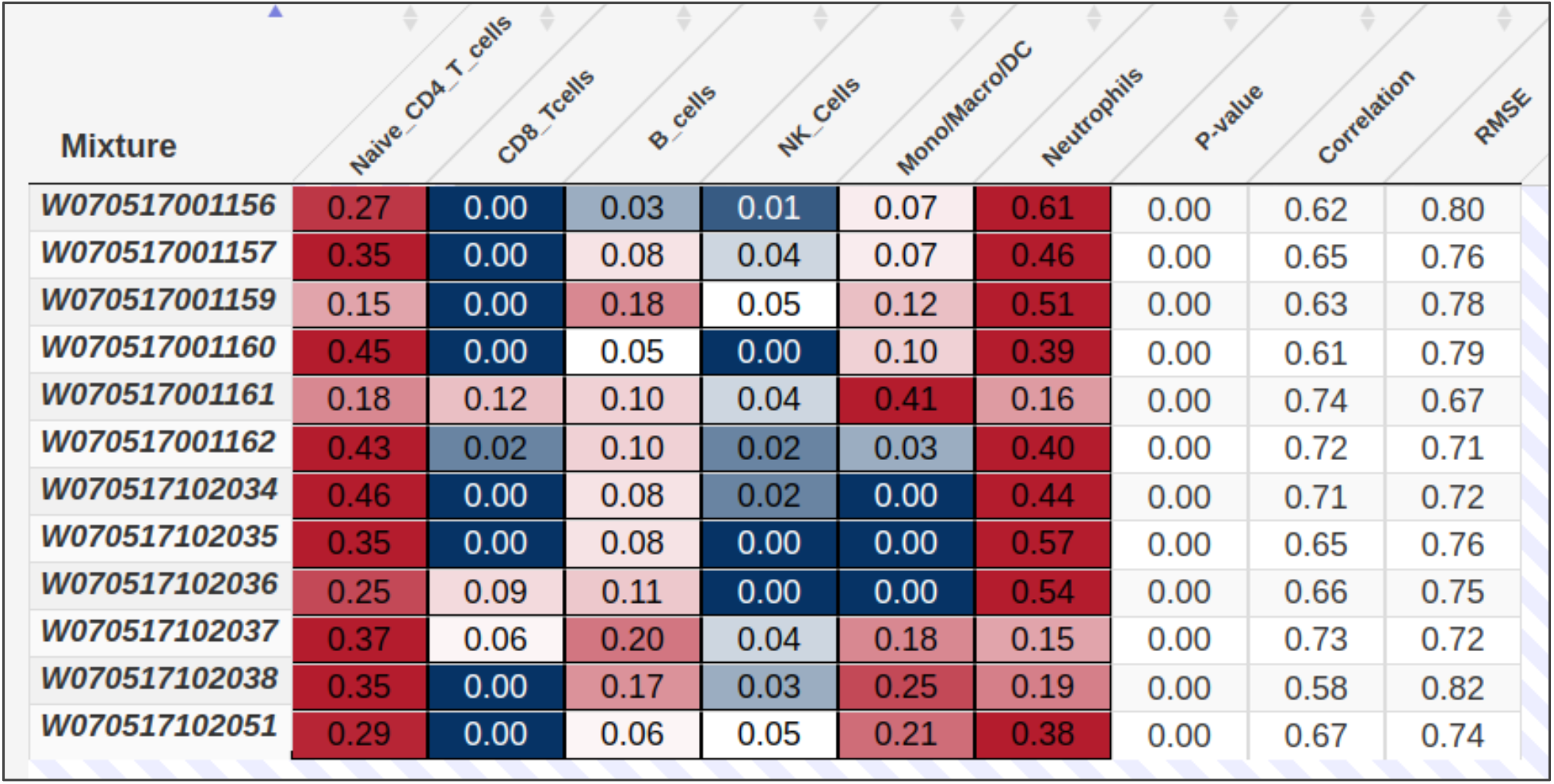
The CIBERSORTx deconvolution results in positive control 1 (whole blood having known proportions of immune cells). The results of the deconvolution by the custom signature matrix were found to be highly correlated with the flow cytometry-derived fractions.

The results of a CIBERSORTx deconvolution analysis were shown in a table, including deconvolution scores for each sample of the mixture in the rows and cell types from the custom signature matrix in the columns. All findings were presented as relative fractions across all cell subsets, normalized to 1. The leftmost column contained the names of the mixtures.

**Table 1.** Flow cytometry derived cell type proportions of lymphocytes in the whole blood (ground truth-positive control 1).

| samples | Neutrophils | Lymphocytes | Monocytes | T cells | T cells CD8 | T cells CD4 | B cells | NK cells |
| --- | --- | --- | --- | --- | --- | --- | --- | --- |
| W070517001156 | 0.796 | 0.105 | 0.099 | 0.066 | 0.015 | 0.041 | 0.01 | 0.021 |
| W070517001157 | 0.66 | 0.259 | 0.081 | 0.193 | 0.048 | 0.147 | 0.039 | 0.031 |
| W070517001159 | 0.77 | 0.148 | 0.082 | 0.044 | 0.008 | 0.035 | 0.052 | 0.058 |
| W070517001160 | 0.65 | 0.244 | 0.106 | 0.21 | 0.032 | 0.19 | 0.024 | 0.025 |
| W070517001161 | 0.537 | 0.281 | 0.183 | 0.223 | 0.142 | 0.072 | 0.032 | 0.07 |
| W070517001162 | 0.489 | 0.433 | 0.078 | 0.28 | 0.082 | 0.193 | 0.059 | 0.1 |
| W070517102034 | 0.615 | 0.321 | 0.064 | 0.202 | 0.057 | 0.146 | 0.069 | 0.027 |
| W070517102035 | 0.776 | 0.148 | 0.077 | 0.118 | 0.04 | 0.066 | 0.03 | 0.015 |
| W070517102036 | 0.6 | 0.283 | 0.118 | 0.191 | 0.077 | 0.107 | 0.054 | 0.038 |
| W070517102037 | 0.46 | 0.45 | 0.09 | 0.267 | 0.116 | 0.156 | 0.115 | 0.069 |
| W070517102038 | 0.598 | 0.326 | 0.076 | 0.207 | 0.057 | 0.156 | 0.082 | 0.037 |
| W070517102051 | 0.692 | 0.169 | 0.138 | 0.099 | 0.032 | 0.073 | 0.028 | 0.057 |

The metrics provided for the results were as follows:

#### P-value

A useful tool for weeding out outcomes with an inadequate “goodness of fit”; to find out if the results of the deconvolution are statistically significant for all cell subtypes.

#### Pearson’s correlation

(R) is the result obtained when the estimated population percentages are compared to the actual population percentages. The predicted cellular fractions and expression profiles from the signature gene matrix were used to make the estimated mixture. It is to be noted that the correlation was calculated for the signature genes only, and not all the genes present in the mixture.

#### RMSE

stands for Root Mean Squared Error. It is the difference amid the actual mixture and the predicted mixture, but only for genes in the signature gene file.

### 3.2. Statistical analysis - Correlation and regression

The concordance between the CIBERSORTx derived proportions and the flow cytometry derived fractions was determined by the Pearson correlation coefficient for each cell type. The B lymphocytes, CD4 T lymphocytes, CD8 T lymphocytes, and neutrophils showed a strong correlation of 0.8 and above, monocytes also showed a strong correlation of 0.7. The NK cells showed a moderately strong correlation of 0.5. The plot shows correlation values between the CIBERSORTx results and the flow cytometry results, for each cell subset.

The overall correlation from the regression analysis was 0.74, and the p-value was below 0.05, which implies that the relationship among the two values was positive and the CIBERSORT results were reliable and in line with the flow cytometry results. The plot shows the regression analysis for all the cell types, and the regression line is shown in blue.

#### Positive control 2 and negative control

The whole blood + purified immune cell subsets (positive control 2), and the neuroblastoma cell lines (negative control) were subjected to the deconvolution, one by one, using the merged custom signature matrix. Both samples were run twice, first in the relative mode to find out relative fractions and later in the “absolute” mode to find out the original absolute immune cell level in every sample for each cell type and also the overall immune content in each sample.

The overall absolute score of immune subsets from the signature matrix in the mixture was high in the whole blood + immune samples but lower or almost negligible in the neuroblastoma cell line sample, as expected.

When compared for individual cell types, the same was observed for both types of samples. The negative control had very low levels for each individual cell type. Noticeably, the positive control asserts the accuracy of the signature matrix. For example, all the NK cell samples showed a very high absolute score for NK cells in the signature matrix. The same is true for all other individual cell types. The plot also demonstrates that the signature gene profiles of T cells (CD4 and CD8) are very similar, as it showed a high content of CD4 T cells even in CD8 samples. This means that CD8 T cells are also in the CD4 group, which means that more sorting or sub clustering is needed, which might be true as the CD4 cluster was the biggest in the single-cell data (4793 cells), from which the signature matrix was generated. Similarly, the monocytes and neutrophils have similar profiles, so they have similar levels in the monocyte sample. This could also mean that the neutrophil cluster also had monocyte cells, which is why it had a high score in the monocyte sample. More sub clustering of the neutrophil cluster is needed to separate the monocytes and find better marker genes that are uniquely found in neutrophil cells.

### 3.3. Deconvolution of TCGA-LUAD data

After validating the signature matrix, we used the custom signature matrix of 14 immune cell subsets to deconvolve the TCGA-LUAD data and find out the content of each type of immune cell in the cancer data. All 598 samples were subjected to the deconvolution process. The results of a CIBERSORTx deconvolution analysis were shown in supplementary files (“LUAD_CIB_rel_fractions” – showing relative score, “LUAD-CIB_ABS_score” – showing absolute scores) including deconvolution scores for each sample of the mixture in the rows and cell types from the custom signature matrix in the columns. All findings were presented as relative fractions across all cell subsets, normalized to 1. The leftmost column contained the names of the mixtures. The metrics provided for the results were P-value, Pearson’s correlation, and RMSE (described in section 3.1)

### 3.4. ESTIMATE Analysis

The ESTIMATE analysis helped in finding out the extent of immune and stromal infiltration within the cancer data. It gave us three types of scores:

- Stromal score (This demonstrates the presence of stroma in tumor tissue),
- Immune score (This indicates the presence of immune cells in the tumor site.), and
- ESTIMATE score (implying the purity of the tumor).

The stromal score ranged from -1822 to 2003, the immune score ranged from -1567 to 2771 and the ESTIMATE score or the tumor purity score, ranged from -3135 to 3975 (Supplementary file “ESTIMATE_LUAD”). The information obtained can be correlated with the patient’s overall survival, which has been done in another paper. It’s the same for the absolute score for the deconvoluted cell types. This information can also be linked to whether or not a certain cell type (determined by deconvolution) is good or bad for the patient’s survival.

CIBERSORTx has its own signature matrix, the “LM22,” which is well validated with computationally generated as well as lab generated data. This LM22 was created using the microarray data. The goal of this study was to utilize the single-cell data to generate a custom signature matrix so that there is precision in terms of signature genes for each cell type, as single-cell data provides high-resolution. Therefore, the LM22 was not utilized for deconvolution in this work.

The signature matrix had to be tested in its entirety, so none of the subtypes could be left out. This is the reason why the subtypes in the custom signature matrix were merged together into major cell types. The merging of the cell types is supported by the heatmap obtained by hierarchical clustering in the previous paper (Verma, 2024), **figure.9 “*Heatmap showing the differential expression of the canonical markers in the custom signature matrix*.*”***. This heatmap establishes the reliability of the merging of cell subtypes. It has the two NK cell subtypes grouped together, all the CD8 subtypes are grouped together, all the B cells subtypes grouped together, the myeloids grouped toghether and neutrophils and naive CD4 T cells are single groups.

**Figure 2.**
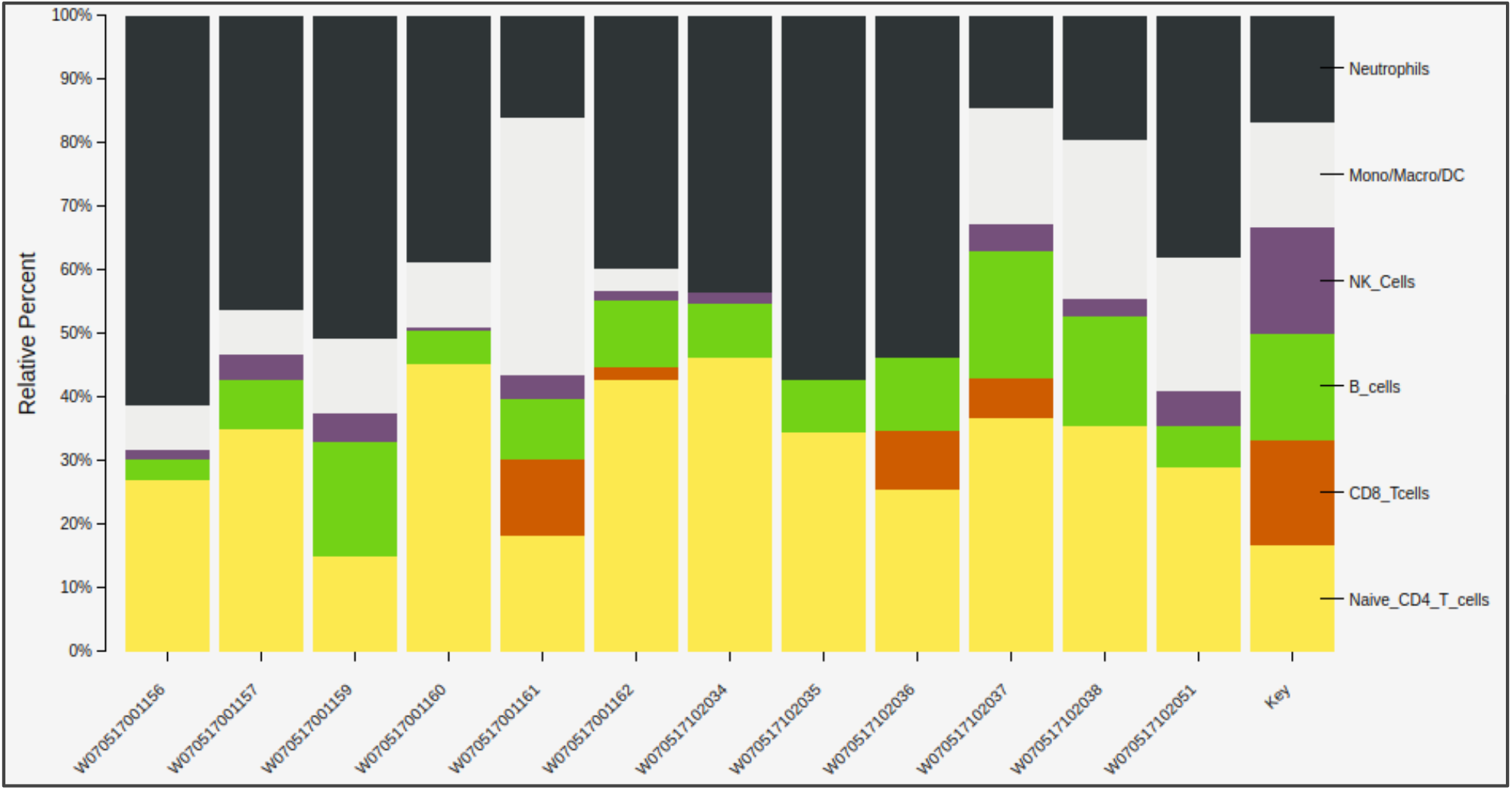
A stacked bar plot depicting the relative percentages of immune cells in each positive control 1 sample.

**Figure 3.**
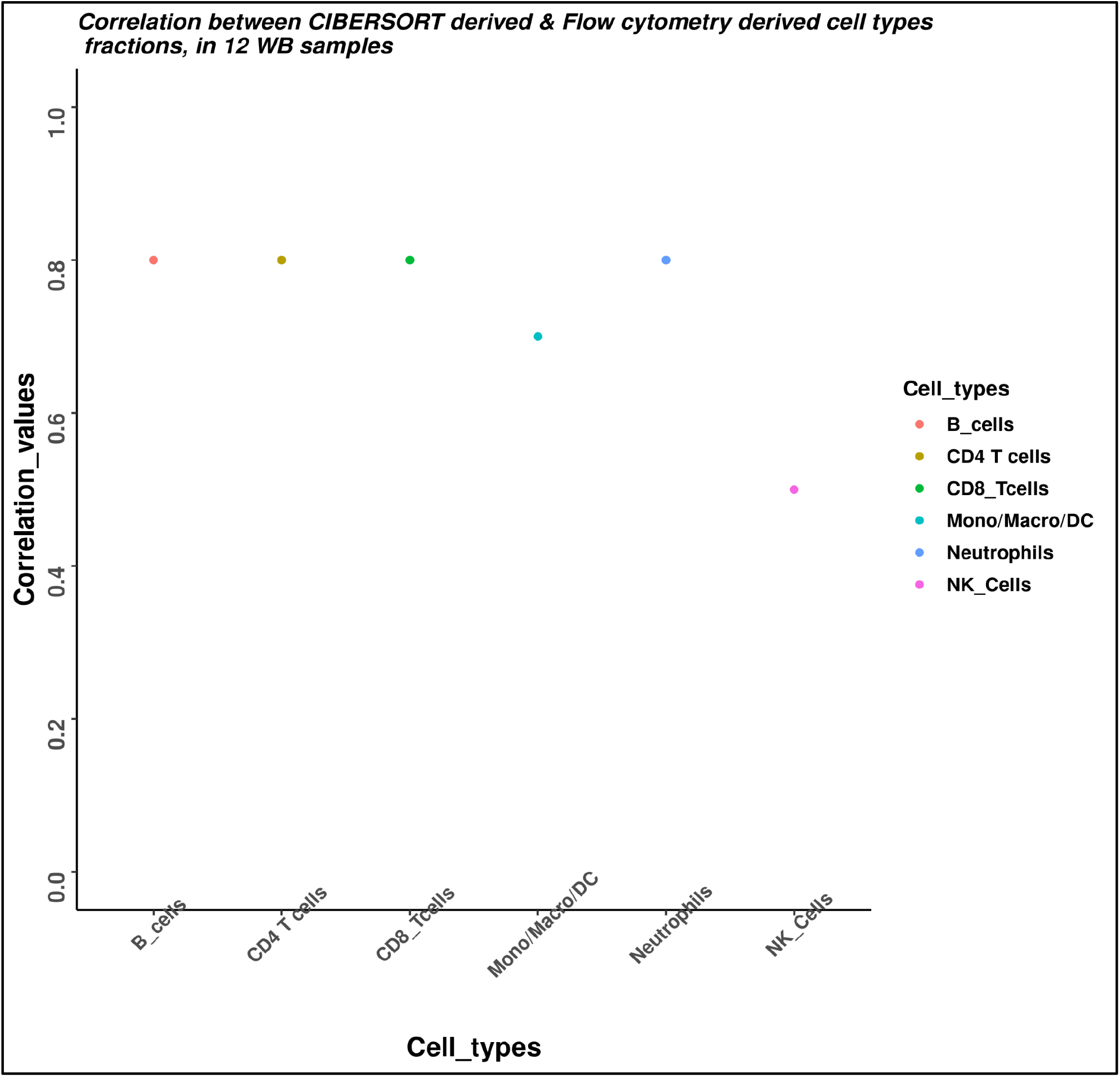
Correlation values obtained when known proportions were compared with CIBERSORTx predicted proportions, for each immune cell type, in positive control 1(Whole blood). All the cell types have a high correlation with the flow cytometry values, except for NK cells, which could be because of the underrepresentation of the data (many zeros), which is why positive control 2 was also tested.

**Figure 4.**
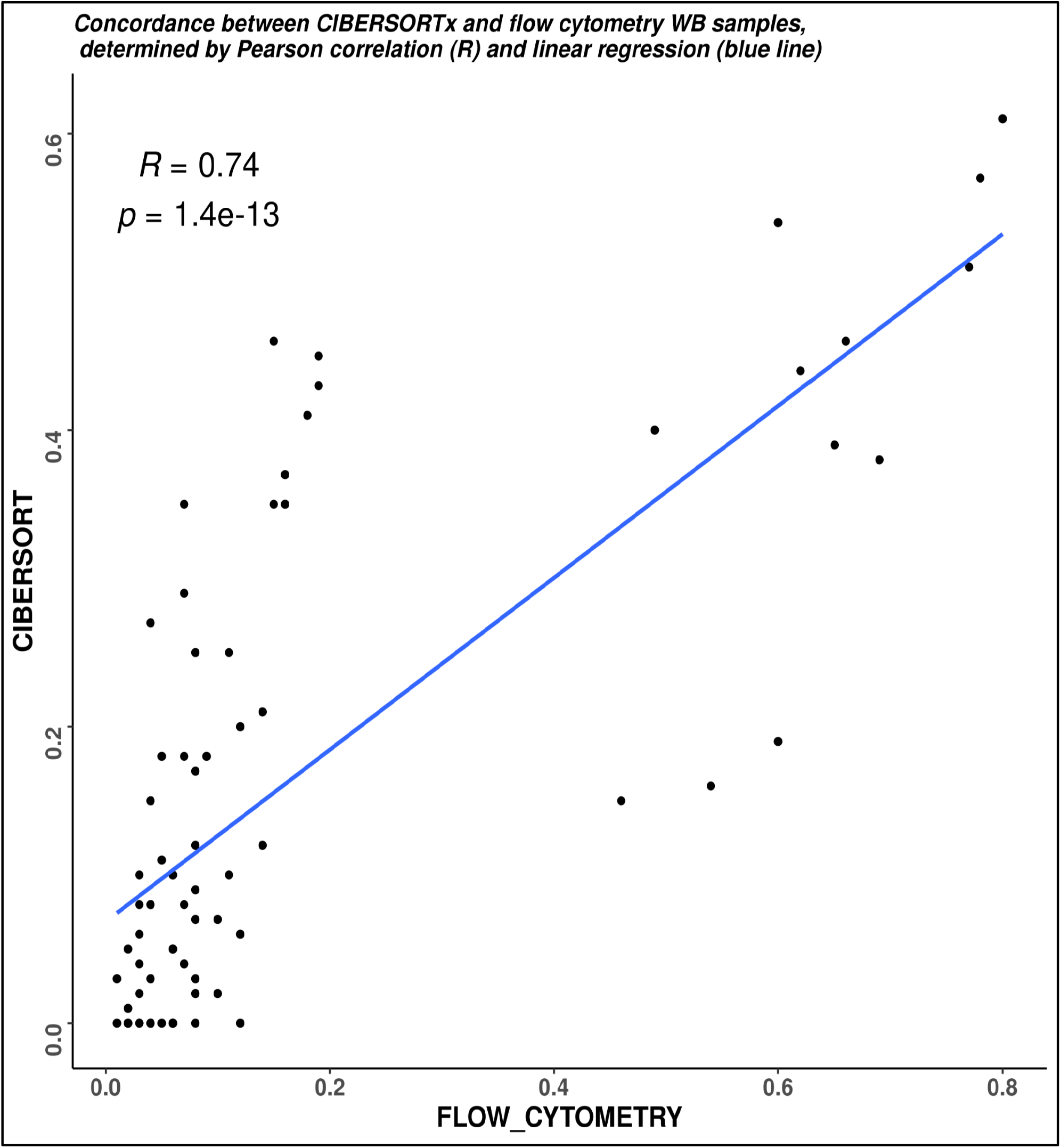
Regression analysis for the CIBERSORTx predicted immune cell proportions and the flow cytometry derived proportions in the whole blood sample for every cell type. Overall, the CIBERSORTx derived fractions were highly correlated with the actual flow-cytometry derived fractions.

**Figure 5.**
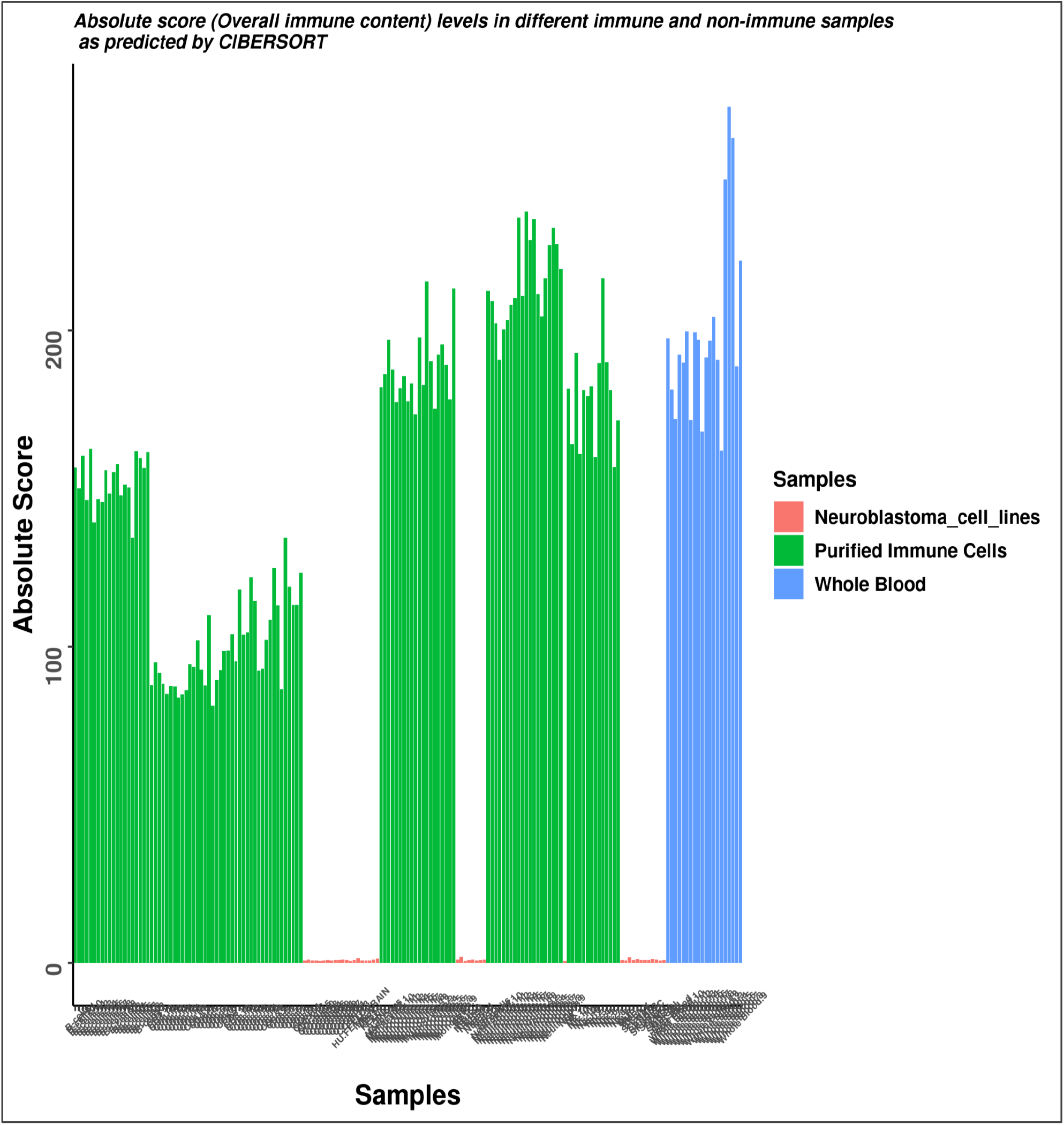
Neuroblastoma Cell lines, whole blood, and purified immune cells’ absolute score (Total immune content) in a barplot. As expected, the positive control has high absolute score levels for overall immune content, contrary to the negative control, which is composed of non-immune or non-hematopoietic samples

**Figure 6.**
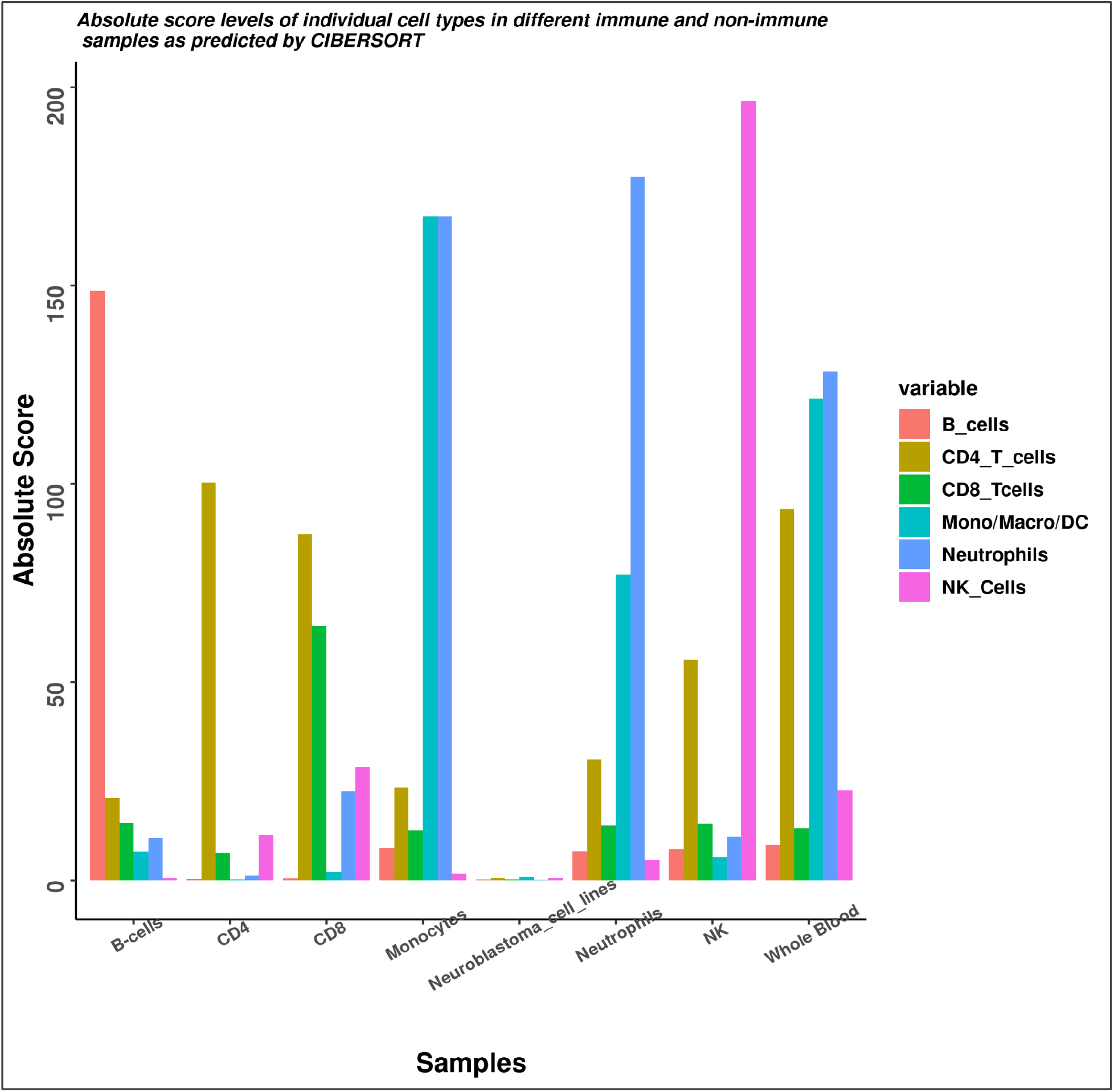
Neuroblastoma Cell lines whole blood and purified immune cells bar plot deconvolution results with the merged 6 cell types custom signature matrix. The negative control has very low levels for each cell type. Noticeably, the positive control asserts the accuracy of the signature matrix. All the NK cell samples show a very high score for NK cells in the signature matrix. The same is true for all other individual cell types.

**Figure 7.**
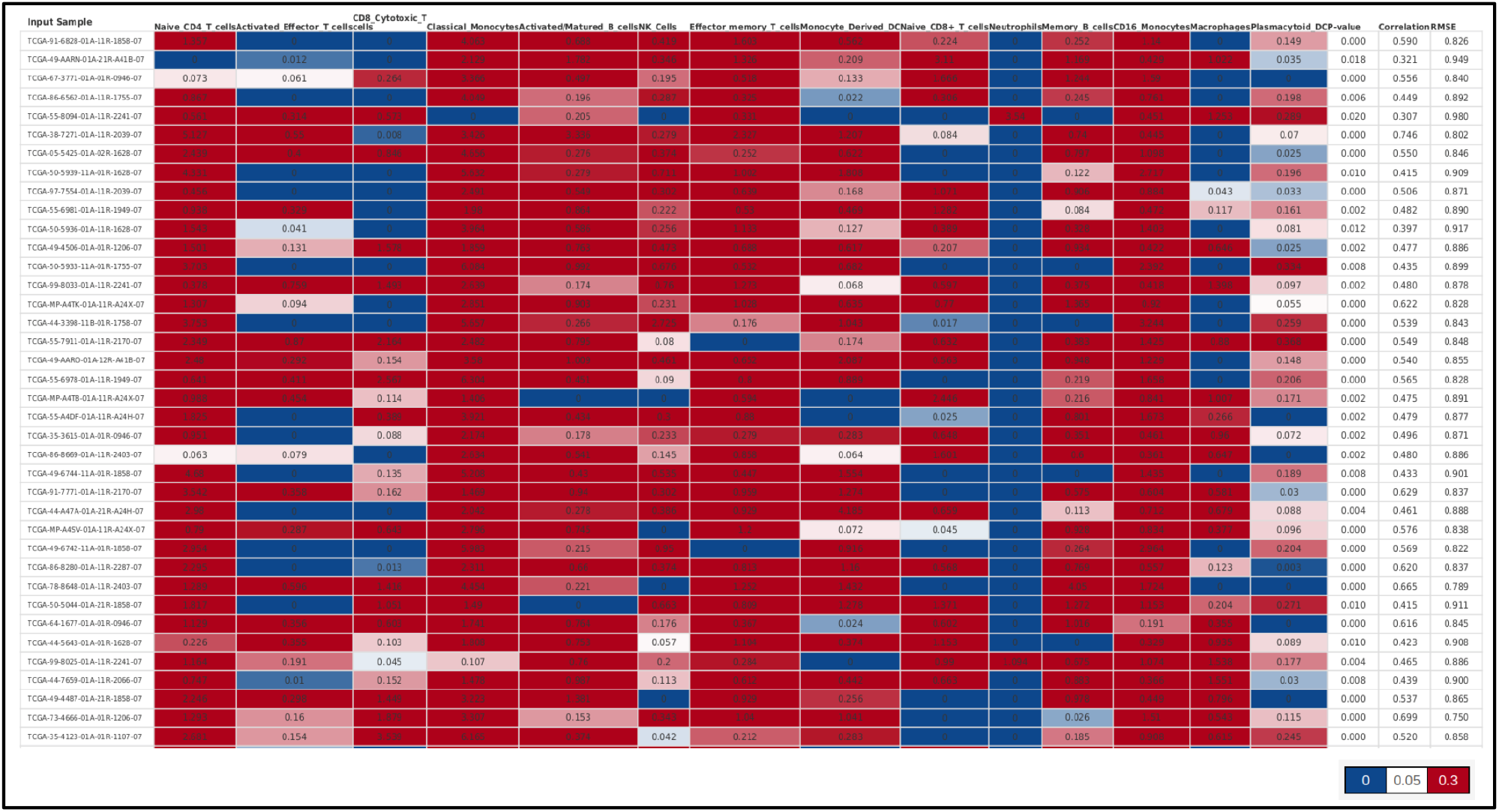
Results of deconvolution of TCGA-LUAD samples using custom signature matrix, by CIBERSORTx (Absolute scores)

**Figure 8.**
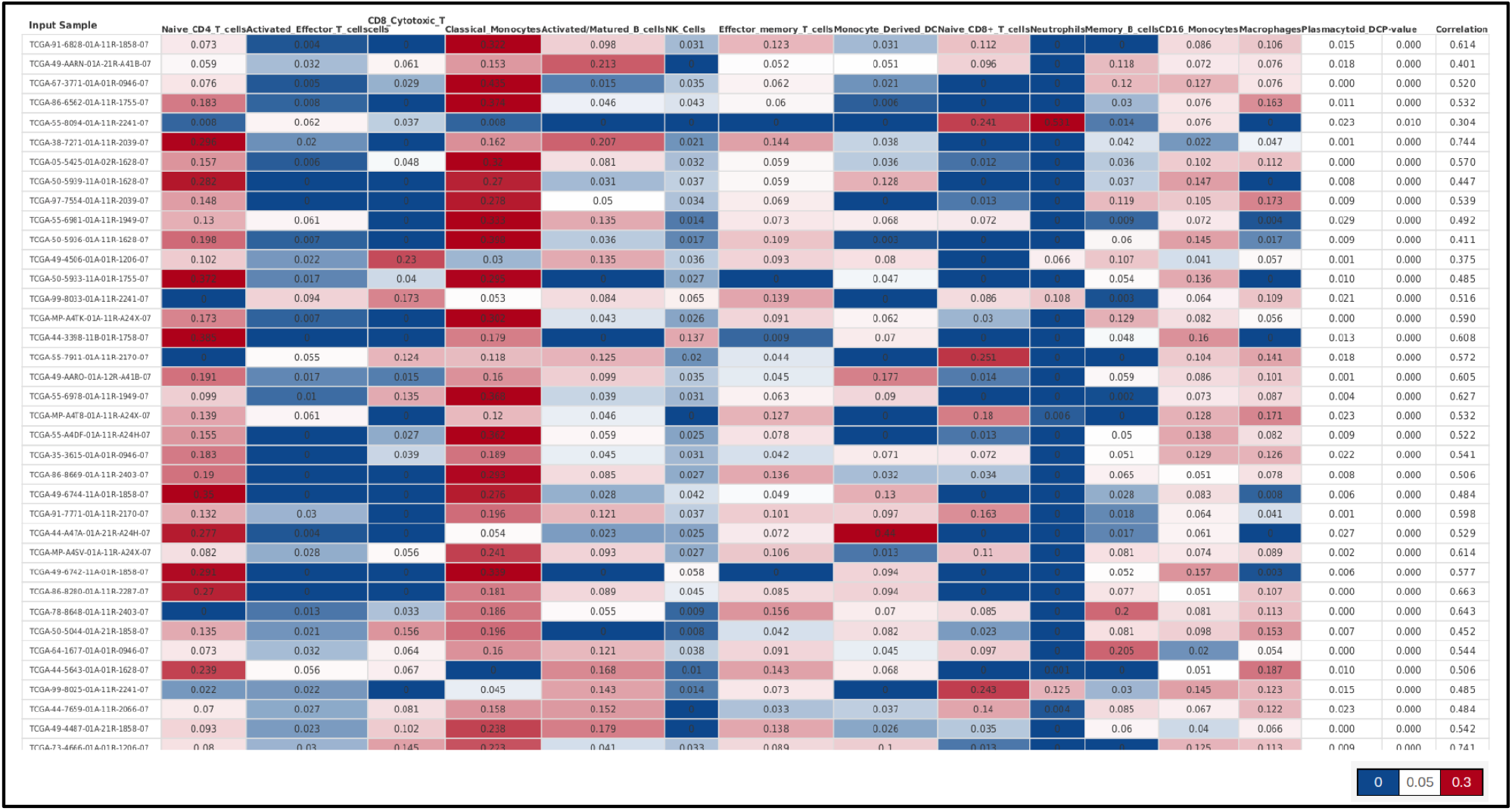
Results of deconvolution of TCGA-LUAD samples using custom signature matrix, by CIBERSORTx (Relative fractions)

The authors of CIBERSORTx suggested methods to validate the custom signature matrix created by the users. The same procedure was followed for validation in this study.

## 4. Conclusion

Validating the custom signature matrix with positive control 1 (flow cytometry determined fractions) gave satisfactory results. The CD8 T cells, CD4 T cells, monocytes, and B cells fractions demonstrated a high positive correlation with the flow cytometry determined fraction. The NK cells demonstrated a moderate correlation with the flow cytometry derived fraction. The reason could be an underrepresentation of the values or the zeros. Overall, the correlation between the two values (CIBERSORT and flow cytometry) was 0.7, and the p-value was below 0.05. This implies that the predicted results were in concordance with the actual flow-cytomtetry-derived results. Using the negative control (neuroblastoma cell lines) and the positive control 2 (whole blood and purified immune cells) to validate the custom signature matrix also gave satisfactory results. As expected, the absolute score for the total immune content was low or negligible in the case of NB cell lines but high in the case of WB and purified immune cells. Also, the absolute score for individual cell types gave highly accurate predictions for each purified sample (e.g., NK cells had the highest score in the NK cell sample, and so on…). We applied this custom signature matrix to TCGA-LUAD data (from TCGA) to determine various immune cell percentages so that we could test whether being high or low in LUAD has some significant effect on the overall survival of the tumor patients. We also applied ESTIMATE, to determine the total immune and stromal content in every sample, as well as the tumor purity, to again, find out its effect on the overall survival of the patients.

## Appendix

### Supplementary files

- LUAD_CIB_rel_fractions : CIBERSORTx deconvolution of TCGA-LUAD data using custom signature matrix (relative fractions)
- LUAD-CIB_ABS_score : CIBERSORTx deconvolution of TCGA-LUAD data using custom signature matrix (absolute score)
- ESTIMATE_LUAD : ESTIMATE results of TCGA-LUAD data

## Abbreviations

DCs: Dendritic cells
ESTIMATE: Estimation of Stromal and Immune Cells in Malignant Tumor Tissues Using Expression Data
FACS: Fluorescence Activated Cell Sorting
FPKM: Fragments Per Kilobase Million
GEO: Gene Expression Omnibus
GEP: Gene expression profiles
LC: Lung Cancer
LM22: Leucocyte Matrix 22
LUAD: Lung Adenocarcinoma
NB cell lines: Neuroblastoma cell lines
NK cells: Natural Killer cells
NOS: Not otherwise specified
NSCLC: non-small cell lung cancer
PBMC: Peripheral Blood Mononuclear Cells
RMSE: Root Mean Square Error
scRNA-seq: Single cell RNA sequencing
ssGSEA: Single-sample gene set enrichment analysis
T cells: T lymphocytes
T-regs: Regulatory T Cells
TCGA: The Cancer Genome Atlas
TIICs: tumor-infiltrating immune cells
TILs: Tumor infiltrating lymphocytes
TMM: trimmed mean of M values
WB: Whole Blood

## Notes

### Competing Interest Statement

The authors have declared no competing interest.

